# Structure-Informed Stress-Testing of the Standard Numbering Scheme for Class B β-Lactamases Using AlphaFold2

**DOI:** 10.64898/2026.10.01.756119

**Authors:** Muhammad Musshif Khan Niazi, Haris Ahmed Khan

**Author notes:** Corresponding author: Muhammad Musshif Khan Niazi.

## Abstract

The standard numbering scheme for class B β-lactamases (metallo-β-lactamases, MBLs), the BBL scheme, was proposed by Galleni et al. in 2001 and updated by Garau et al. in 2004 from the small number of MBL structures then solved. Hundreds of additional class B sequences have since been catalogued, but most have never been structurally solved, so whether BBL correspondences hold across this larger diversity has not been directly tested. Following the precedent set for the sibling class D enzymes, we used ColabFold (AlphaFold2/MMseqs2) to model 87 class B families across subclasses B1, B2, and B3 lacking an experimental structure, after validating the pipeline against an eight-enzyme benchmark spanning all three subclasses (RMSD to experimental structures 0.35–0.46 Å). Applying BBL zinc-ligand numbering to the 87 models via paired sequence- and structure-based residue-transfer methods, the canonical zinc-coordination motif transferred cleanly in the large majority of families, and 11 of the 16 sequence-level exceptions were accounted for by a literature-supported biological cause (a subclass-appropriate substitution or a previously proposed, unconfirmed fourth subgroup), while two motif-retaining families showed predicted local structural divergence at the active-site loop, flagged by structural superposition, that could not be resolved to exact zinc-ligand numbering and warrants direct experimental follow-up. These results show that the BBL scheme, formulated before metagenomic sequencing or deep-learning structure prediction existed, generalizes robustly into the structural space these technologies have since opened, and identify a small set of variants whose active sites merit experimental attention.

**IMPORTANCE:** Metallo-β-lactamases inactivate carbapenems, antibiotics of last resort for multidrug-resistant Gram-negative infections, and new MBL variants are discovered faster than they can be structurally characterized. The BBL numbering scheme underlies how researchers compare active-site residues across this diverse family, but it was built from only a handful of solved structures and has not been re-examined against the much larger sequence diversity now known. We show that validated AlphaFold2 prediction can test whether a numbering scheme built on a small structural sample still holds for unsolved enzymes, and that most of class B is well described by the existing scheme. This approach also flags enzymes whose active site appears diverged from the canonical arrangement, prioritizing them for structural work and giving surveillance and inhibitor-design efforts a validated numbering reference covering far more class B diversity than the eight enzymes on which the scheme was built.

## INTRODUCTION

β-Lactamases are the single most important mechanism of bacterial resistance to β-lactam antibiotics and are classified into four molecular classes (A–D) on the basis of amino acid sequence (1). Classes A, C, and D use an active-site serine nucleophile for catalysis, whereas class B enzymes — the metallo-β-lactamases (MBLs) — are zinc-dependent hydrolases with no sequence or structural relationship to the serine enzymes (2–4). MBLs are of particular clinical concern because they efficiently hydrolyze carbapenems, the β-lactam antibiotics typically reserved for infections resistant to other agents, and because no MBL inhibitor is yet in routine clinical use (2, 5). On the basis of sequence relationships, class B enzymes are divided into three subclasses — B1, B2, and B3 — which differ in oligomeric state, zinc stoichiometry and coordination chemistry, and substrate range, despite sharing a common overall αβ/βα metallo-hydrolase fold (6, 7).

Because class B enzymes from different subclasses share as little as ∼20% pairwise sequence identity, comparing functionally equivalent residues across the family using each enzyme’s own native numbering is unreliable and has historically produced inconsistent descriptions of the same catalytic positions in the literature (6). To resolve this, Galleni et al. proposed a standard numbering scheme for class B β-lactamases in 2001 — the BBL scheme — anchored to the sequence of the Stenotrophomonas maltophilia L1 enzyme and built from the small number of class B crystal structures solved at that time (BcII, CcrA, and IMP-1 in subclass B1, and L1 in subclass B3) (6). Garau et al. updated this scheme in 2004 once a subclass B2 structure (CphA) became available, using a structure-based multiple alignment (via the TOP/MAPS algorithm) of all eight class B structures solved by that date to refine the numbering and to identify fourteen sequence fragments of structurally conserved position spanning the whole family (7). Under the BBL scheme, the canonical zinc-coordinating residues are His116, His118, and His196 (Zn1) together with Asp120, Cys221, and His263 (Zn2) in subclass B1; the same positions but with Asn116 replacing His116 in subclass B2; and the same positions but with His121 replacing Cys221 — contributed from a structurally distinct part of the polypeptide chain — in subclass B3, where His/Gln116 is also tolerated at the first Zn1 position (6, 7). These correspondences, along with the observation that a loop closing the B1 active site (BBL residues 61–65) is present in subclass B1 but absent from the solved B3 and (probably) B2 structures, remain the reference framework cited whenever a newly described class B enzyme’s active site is compared with the rest of the family.

The BBL scheme’s numbering has proved durable — Garau et al. themselves noted that essentially all of the important residue correspondences were unchanged between the 2001 and 2004 versions, with revisions confined mainly to the N- and C-termini (7) — but it was necessarily built from a very small structural sample: eight solved structures as of 2004, versus the many hundreds of class B sequences now catalogued in resources such as the Beta-Lactamase DataBase (BLDB) (8). The great majority of named class B families still have no experimentally determined structure at all, so whether the BBL correspondences continue to hold for this much larger and more diverse body of sequence — much of it from environmental, agricultural, or otherwise atypical bacterial sources rather than the clinical Enterobacterales, Pseudomonas, and Acinetobacter isolates from which the original reference structures were drawn — has not been directly tested.

Deep-learning structure prediction offers a way to close this gap without waiting for crystallography or cryo-EM. Stasyuk and Smith recently demonstrated this approach for the sibling class D (OXA-type) β-lactamases: after building a structure-based numbering and secondary-structure scheme from 25 solved class D structures, they used AlphaFold2 (via the ColabFold implementation) to predict structures for four class D enzymes representing categories of increasing structural novelty — a member of a family with a solved relative, a member of a family with no solved representative, an orphan enzyme with no family, and a novel enzyme from a Gram-positive organism not previously known to carry class D β-lactamases — and showed that their numbering scheme transferred cleanly onto all four AlphaFold2 models (9). This established both a template methodology and, more specifically, that a residue-numbering scheme built from a small experimentally solved subset of a β-lactamase class can be validated and extended into much larger, structurally unsampled sequence space using AlphaFold2 predictions.

We apply the same logic to class B (workflow in Figure 5). We first validate a ColabFold-based structure-prediction pipeline against an eight-enzyme experimental benchmark spanning all three class B subclasses, then use the validated pipeline to predict structures for 87 class B families that currently have no solved structure of any kind, drawn from across subclasses B1, B2, and B3. We use these predicted structures — via paired sequence-based and structure-based residue-transfer methods — to test whether the BBL zinc-ligand numbering and the subclass-diagnostic active-site loop continue to hold across this structurally unsampled diversity, and we characterize the specific exceptions we find.

We note that Oelschlaeger et al. have separately identified the same underlying problem — that class B numbering and naming conventions have not kept pace with the expansion of known sequence diversity — and describe an in-development, sequence-based renumbering tool built on profile hidden Markov models rather than on structure prediction (10). Because that approach operates entirely on sequence and does not generate or evaluate three-dimensional models, it cannot by itself detect the kind of local active-site structural divergence we identify below in two of the 87 families examined here; we discuss this distinction further below.

## RESULTS

### Dataset construction

We began from the full class B catalogue in the Beta-Lactamase DataBase (BLDB) (8), comprising 1,021 individually named variants across 129 family–subclass entries (128 unique family names; PEDO is listed under both B1 and B3) (subclass B1: 686 variants, 69 families; B2: 25 variants, 5 families; B3: 310 variants, 55 families). Families for which BLDB listed no solved PDB structure for any member variant were flagged as candidate targets (n = 96 family–subclass entries; 95 unique families) and cross-checked in three independent passes: (i) against the 37 protein families of the validation set of enzymes with confirmed solved structures (described below), which identified three BLDB annotation errors (CfiA, MEMA1, and RM3 in fact have solved structures and were removed from the candidate list); (ii) by direct RCSB PDB and literature search for a spot-check subsample, which confirmed several candidates as genuinely unsolved and two (VIM, LRA-12) as correctly excluded solved positive controls; and (iii) by an independent literature pass across the remaining candidates, which identified three further families (AFM, CIM2, CJO) that had been solved since the BLDB entry was last updated (CIM2 and CJO corresponding to PDB entries 9GX8 and 9GX9, respectively) and removed them. A further family, CAU, was found by direct sequence comparison to be 98% identical to a deposited structure (PDB 8U00) and was likewise reclassified as solved and removed. One additional family, SZM, was excluded because no individual GenBank protein accession exists for it (only a BioProject-level deposition), precluding structure prediction from an individual sequence. This process yielded a final target set of 87 class B families with no solved structure, spanning subclasses B1 (n = 46), B2 (n = 3), and B3 (n = 38) by BLDB subclass assignment (PEDO, which BLDB lists under both B1 [one variant] and B3 [two variants], was counted as B3; the 96 candidate entries, less the duplicate PEDO entry and the eight families removed above, give 87).

A parallel validation set of enzymes with solved structures was assembled from the same BLDB catalogue (60 unique enzymes (deduplicated by protein name) representing 590 individual PDB entries across 37 protein families); sequence retrieval yielded 59 records; MEMA1-1 was excluded because only a fosmid-level BLAST-relative accession, and not an individually deposited sequence for the protein itself, could be identified in either the primary characterization or its supplementary data. On audit, one record (NDM-1, A0A0A7Y424) proved to be a 264-residue fragment identical to part of the full-length NDM-1 record (C7C422) and was removed, leaving 58 unique sequences. The VIM-1 sequence (Q9XAY4) was then added directly, giving 59 sequences. One retained record (B8QIQ9, labelled only blaVIM) differs from VIM-2 by a single residue, which is why 59 sequences represent 58 enzymes. MEMA1-1 and NDM-4 are therefore the two enzymes without a sequence of their own. Because BLDB cross-lists UniProt codes between named enzymes, NDM-4 is not represented by a sequence of its own (its cross-listed accession corresponds to NDM-1); the set therefore contains independently sequenced representatives of 58 of the 60 enzymes. From this validation set we further selected an eight-enzyme historical benchmark subset — BcII, CcrA, IMP-1, VIM-2, and BlaB (subclass B1), CphA (subclass B2), and L1 and FEZ-1 (subclass B3) — corresponding to the eight structures used in the original 2004 BBL update (7), for use as the primary pipeline-validation and structural-anchoring reference set throughout this study.

### Signal peptide removal

All 87 target-set sequences were screened for N-terminal signal peptides using DeepSig (11). DeepSig identified and removed a signal peptide in 69 of 87 sequences directly. Of the remaining 18 sequences called "no signal peptide," a Kyte–Doolittle hydrophobicity scan of the N-terminal region flagged the majority as nonetheless possessing a hydrophobic-core signature consistent with an undetected leader sequence; all 18 were therefore submitted to SignalP 6.0 (organism setting "Other," slow/sensitive mode) as an independent second call (12). This second pass identified and trimmed a signal peptide at a specific, high-confidence cleavage site (probability ≥ 0.70) in 13 of the 18 sequences, two of which (EFM, BMHC) were identified as lipoproteins (SignalP class "LIPO") whose mature sequence correctly begins at the lipidated cysteine. One of the 13 (MSI) required correction of its input sequence: its source record carried an N-terminal extension ahead of an in-frame internal methionine (position 53), and SignalP applied from that methionine identified a leader (probability 0.999) cleaved after residue 22 of the submitted sequence, so that the mature sequence begins at residue 75 of the source record (267 residues). The remaining five sequences (PAN, SFB, SLB, BB32, and AHM) were retained at full length: no cleavage site reached the probability threshold under either method, and three of them (PAN, SFB, and SLB) were genuinely ambiguous even under SignalP (posterior probabilities of approximately 0.50 for the signal-peptide class). These were deliberately not trimmed on a low-confidence call, and this is stated as an explicit limitation of the mature-sequence set (see Discussion). The final corrected mature-sequence set comprised 87 unique sequences ranging from 217 to 372 residues.

### Pipeline validation against the eight-enzyme experimental benchmark

Prior to prediction of the 87 target-set structures, the ColabFold structure-prediction pipeline (13, 14) was validated against the eight-enzyme historical benchmark set for which high-resolution experimental structures are available. All eight enzymes were folded using identical settings (ColabFold v1.6.3; AlphaFold2-ptm models; no structural templates; MMseqs2 UniRef+environmental paired/unpaired MSA; three recycles; top-ranked model relaxed with Amber) and the resulting top-ranked predicted models were superimposed on their corresponding experimental structures in PyMOL. All eight alignments gave a root-mean-square deviation (RMSD) below 0.5 Å (range 0.351–0.457 Å; Table 2), well under the acceptability threshold of 1.5 Å that we set before the comparison was run, and no individual enzyme was an outlier. Because the benchmark sequences were folded at full (precursor) length rather than as signal-peptide-trimmed mature sequences, we additionally ran all eight through SignalP 6.0 and confirmed that each carries a genuine, high-confidence signal peptide (posterior probability 0.975–0.982; Table 3); the predicted models’ overall mean per-residue confidence (pLDDT) was correspondingly high (90.8–94.1) but showed a marked, enzyme-specific drop when restricted to the first 30 residues (43.4–71.8; Table 3), consistent with the expected low-confidence disorder of an unresolved, uncleaved leader peptide and independently corroborating the SignalP calls via an entirely different (structure-confidence-based, rather than sequence-based) method. Because residues in a predicted model with no counterpart in the resolved experimental chain are excluded from the aligned atom set by the alignment algorithm rather than being penalized, the presence of this extra unmatched leader sequence does not inflate the reported RMSD values, which therefore reflect the accuracy of the core, catalytically relevant fold specifically.

**Table 1.** BBL consensus numbering of the canonical zinc-ligand positions by subclass. (**6, 7**).

| Subclass | Zn1 ligand 1 | Zn1 ligand 2 | Zn1 ligand 3 | Zn2 ligand 1 | Zn2 ligand 2 | Zn2 ligand 3 |
| --- | --- | --- | --- | --- | --- | --- |
| B1 | His116 | His118 | His196 | Asp120 | Cys221 | His263 |
| B2 | Asn116 | His118 | His196 | Asp120 | Cys221 | His263 |
| B3 | His/Gln116 | His118 | His196 | Asp120 | His121 | His263 |

**Table 2.** Structural validation of the ColabFold pipeline against the eight-enzyme historical benchmark set. RMSD and aligned-atom counts are all-atom values from PyMOL’s align command with the default five cycles of outlier rejection (see Materials and Methods); the aligned-atom count is therefore substantially smaller than each model’s total atom count by design, not by data loss.

| Enzyme | Subclass | Reference PDB | RMSD (Å) | Aligned atoms |
| --- | --- | --- | --- | --- |
| BcII | B1 | 1BC2 | 0.405 | 1395/4025 |
| CcrA | B1 | 1ZNB | 0.364 | 1461/3817 |
| IMP-1 | B1 | 1DDK (apo) | 0.457 | 1370/3862 |
| VIM-2 | B1 | 1KO3 | 0.387 | 1357/3973 |
| BlaB | B1 | 1M2X | 0.410 | 1398/3963 |
| CphA | B2 | 1X8G (native) | 0.415 | 1461/3958 |
| L1 | B3 | 1SML (chain A) | 0.400 | 1573/4311 |
| FEZ-1 | B3 | 5WCK (supersedes 1K07) | 0.351 | 1715/4433 |

**Table 3.** Fold confidence and signal-peptide status of the eight-enzyme historical benchmark models.

| Enzyme | Length (aa) | Mean pLDDT | pTM | First-30-residue pLDDT | SignalP cleavage site | SignalP probability |
| --- | --- | --- | --- | --- | --- | --- |
| BcII | 257 | 90.81 | 0.84 | 44.34 | 30–31 | 0.977 |
| BlaB | 249 | 92.50 | 0.86 | 57.08 | 22–23 | 0.976 |
| CcrA | 249 | 94.14 | 0.89 | 68.69 | 18–19 | 0.976 |
| CphA | 254 | 92.58 | 0.88 | 54.35 | 27–28 | 0.979 |
| FEZ-1 | 282 | 94.11 | 0.89 | 60.78 | 20–21 | 0.975 |
| IMP-1 | 246 | 93.87 | 0.87 | 71.82 | 18–19 | 0.982 |
| L1 | 290 | 94.05 | 0.89 | 61.55 | 21–22 | 0.978 |
| VIM-2 | 266 | 90.91 | 0.86 | 43.38 | 26–27 | 0.976 |

### Structure prediction quality across the 87-family target set

All 87 target-set sequences were folded under the same ColabFold settings used for benchmark validation. Overall fold confidence was high across the set (mean pLDDT 95.8, mean predicted TM-score [pTM] 0.92; Figure 1A), with the large majority of families exceeding a pLDDT of 90. One family (BB32) was the lowest-confidence model, with a mean pLDDT of 81.3 and pTM of 0.70, about eight pLDDT points below the next-lowest family (AHM, 89.3); BB32’s reduced confidence is concentrated in specific chain segments and is discussed below together with its active-site divergence.

**Figure 1.**
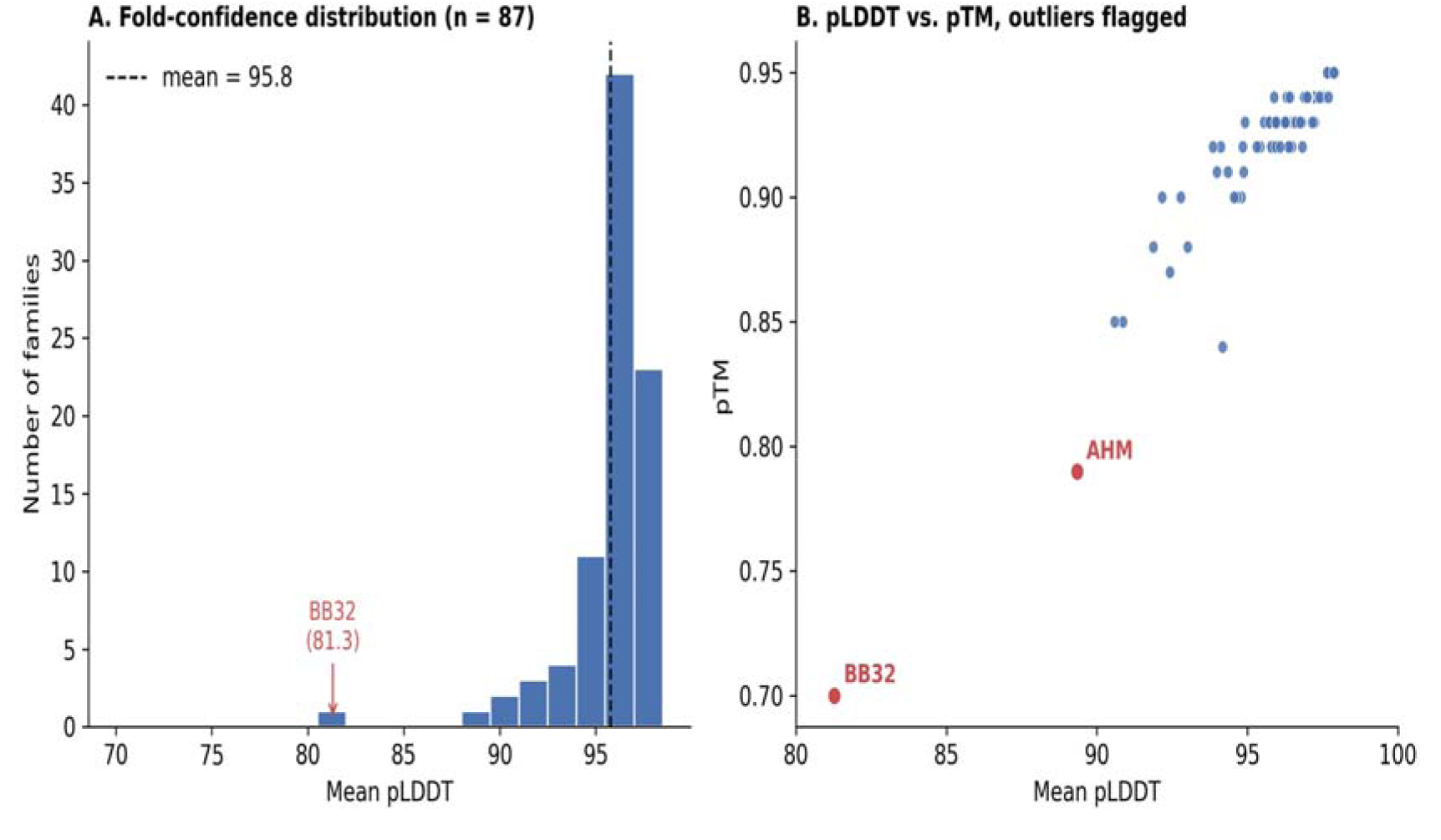
Fold-confidence distribution across the 87-family target set. (A) Distribution of mean per-residue pLDDT across all 87 predicted structures (dashed line, dataset mean = 95.8). The lowest-confidence family, BB32 (mean pLDDT 81.3), is indicated. (B) Mean pLDDT plotted against predicted TM-score (pTM) for all 87 families, with the two lowest-confidence outliers (BB32, AHM) labeled in red.

A signal-peptide tail-confidence check — comparing each family’s overall mean pLDDT to its first-30-residue mean — identified the largest confidence drops in a small number of families (Figure 2), four of which (designated SFB, BB32, SLB and PAN) had been called "no signal peptide" by DeepSig despite showing among the largest such drops in the entire set; this pattern is noted as a candidate signal-peptide-detection miss worth targeted follow-up, though it does not alter the mature-sequence numbering analysis reported below, since, for SFB, SLB and PAN, catalytic-domain confidence away from the N-terminus was unaffected (BB32 is discussed separately below). A per-residue confidence comparison of BB32, SFB, and a representative high-confidence family (Figure 3) illustrates that BB32’s reduced confidence is concentrated in specific chain segments (discussed further below in the context of its active-site divergence), whereas SFB’s is confined largely to the N-terminal tail.

**Figure 2.**
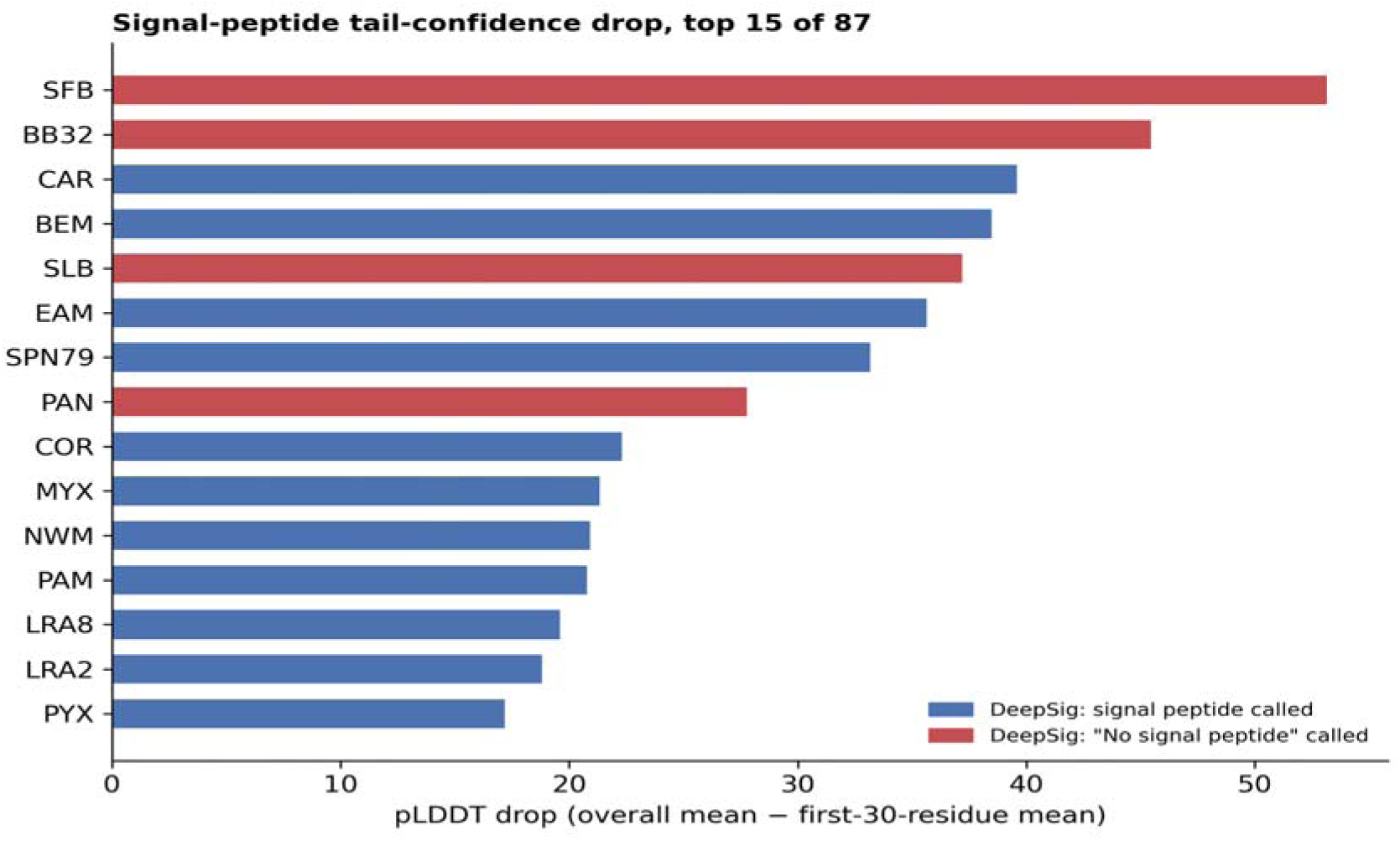
Signal-peptide tail-confidence drop for the 15 target-set families showing the largest difference between overall mean pLDDT and first-30-residue mean pLDDT. Bars are colored by whether DeepSig called a signal peptide (blue) or no signal peptide (red) for that family; four families with a large tail-confidence drop (SFB, BB32, SLB, PAN) were called "no signal peptide" by DeepSig; of these, SFB, SLB, and PAN remained ambiguous under SignalP and are flagged as candidate signal-peptide-detection misses for future follow-up.

**Figure 3.**
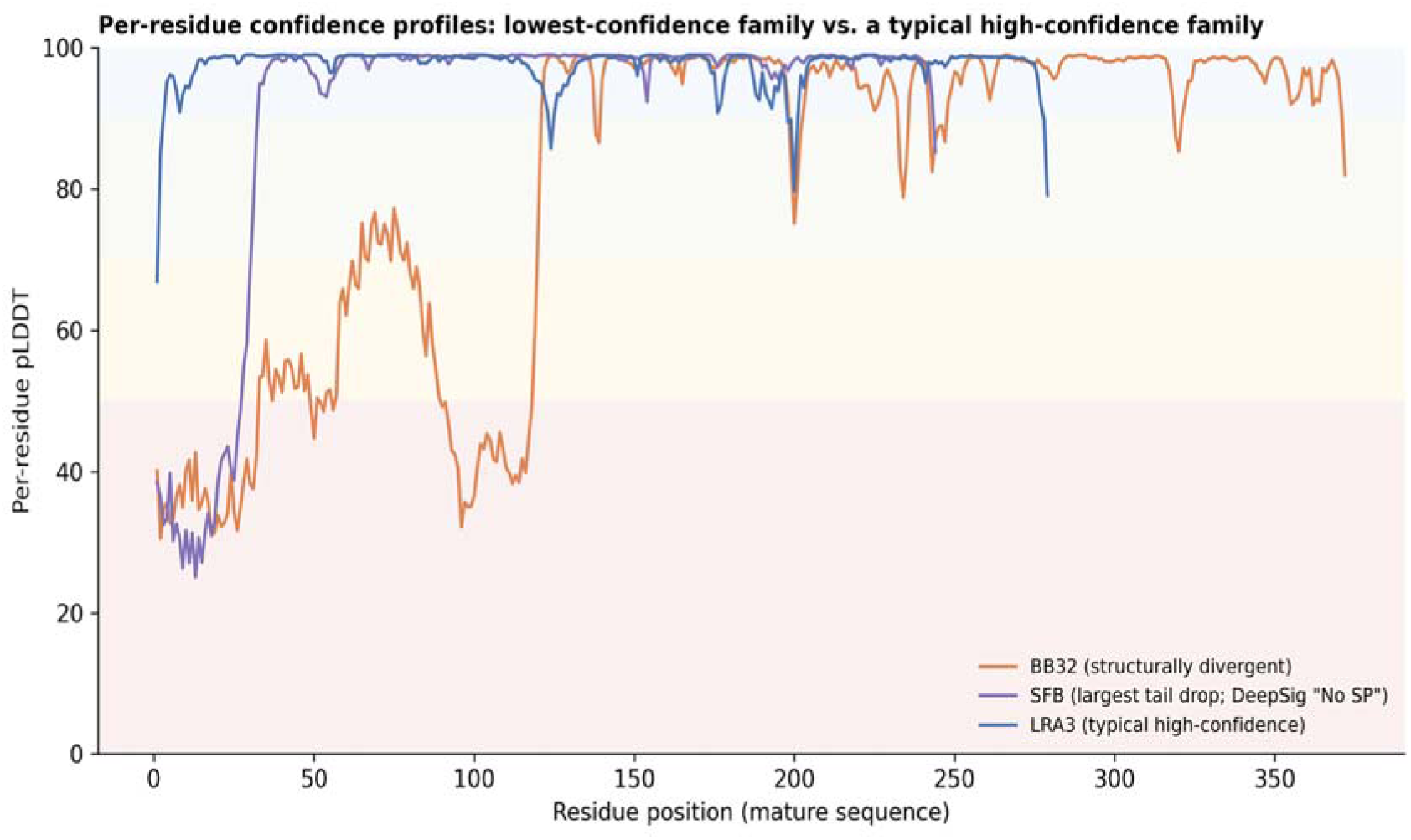
Per-residue confidence profiles for the lowest-confidence family (BB32), the family with the largest signal-peptide tail-confidence drop (SFB), and a representative typical high-confidence family (LRA3), plotted against mature-sequence residue position.

### The BBL zinc-ligand motif is conserved across the great majority of the target set, and 11 of the 16 exceptions have an explained biological cause

As a first, sequence-level test of the BBL numbering scheme’s applicability to the target set, all 87 mature sequences were scanned for the canonical zinc-ligand motif spanning the Zn1 di-histidine (or subclass-appropriate substitute) pair and the Zn2 aspartate (HxHxD, or the subclass-appropriate variants Q/N-x-H-x-D and H-x-D-x-D-H), calibrated against the known mature-sequence positions of the historical benchmark enzymes. Seventy-one of 87 families (82%) showed an exact match to the expected motif. Of the remaining 16 families, literature and BLDB cross-referencing resolved 11 to a specific, citable biological cause (Table 4): three families (PFM, CVI, YEM) were confirmed as genuine subclass B2 members whose naturally divergent Zn1 asparagine (in place of histidine) is the expected, defining B2 substitution rather than an anomaly; seven families (LRA17, LRA19, PEDO, ESP, CPS, PLN, SIQ) were confirmed as subclass B3 members carrying the documented glutamine-for-histidine substitution at the first Zn1 position that is permitted within that subclass; and one family (SPR) showed a genuinely distinct active-site arrangement matching a previously proposed, though not formally established, fourth ("B4") class B subgroup (15), together with its reported close homolog CSA-1. Sixteen families in total initially lacked an exact canonical-motif match; the eleven listed above were resolved to a documented subclass-level biological cause. The remaining five were not among these eleven and were not explained by a subclass-level substitution. Two of them (SER and SSE, subclass B3 by BLDB) match neither the HxHxD nor the Q/N-x-H-x-D pattern (SER carries Arg and SSE carries Glu at a first-Zn1-region position when aligned to L1), so they are reported as genuine motif deviations rather than Gln116 variants. The other three (VMH, AHM, and CrxA — discussed further below) independently overlapped with families that a wholly separate, structure-based analysis (described next) had already flagged as difficult to anchor by simple sequence alignment, supporting the sequence-motif screen as a genuine low-cost triage signal for structurally atypical active sites rather than a source of false positives. Separately, the sequence-alignment screen flagged one exact-HxHxD family, BB11, as carrying Asp rather than His at the BBL 196-equivalent Zn1 position, which we report as an additional candidate deviation not detected by the motif screen.

**Table 4.**
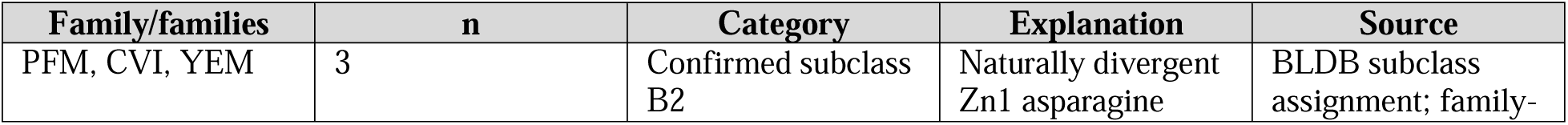
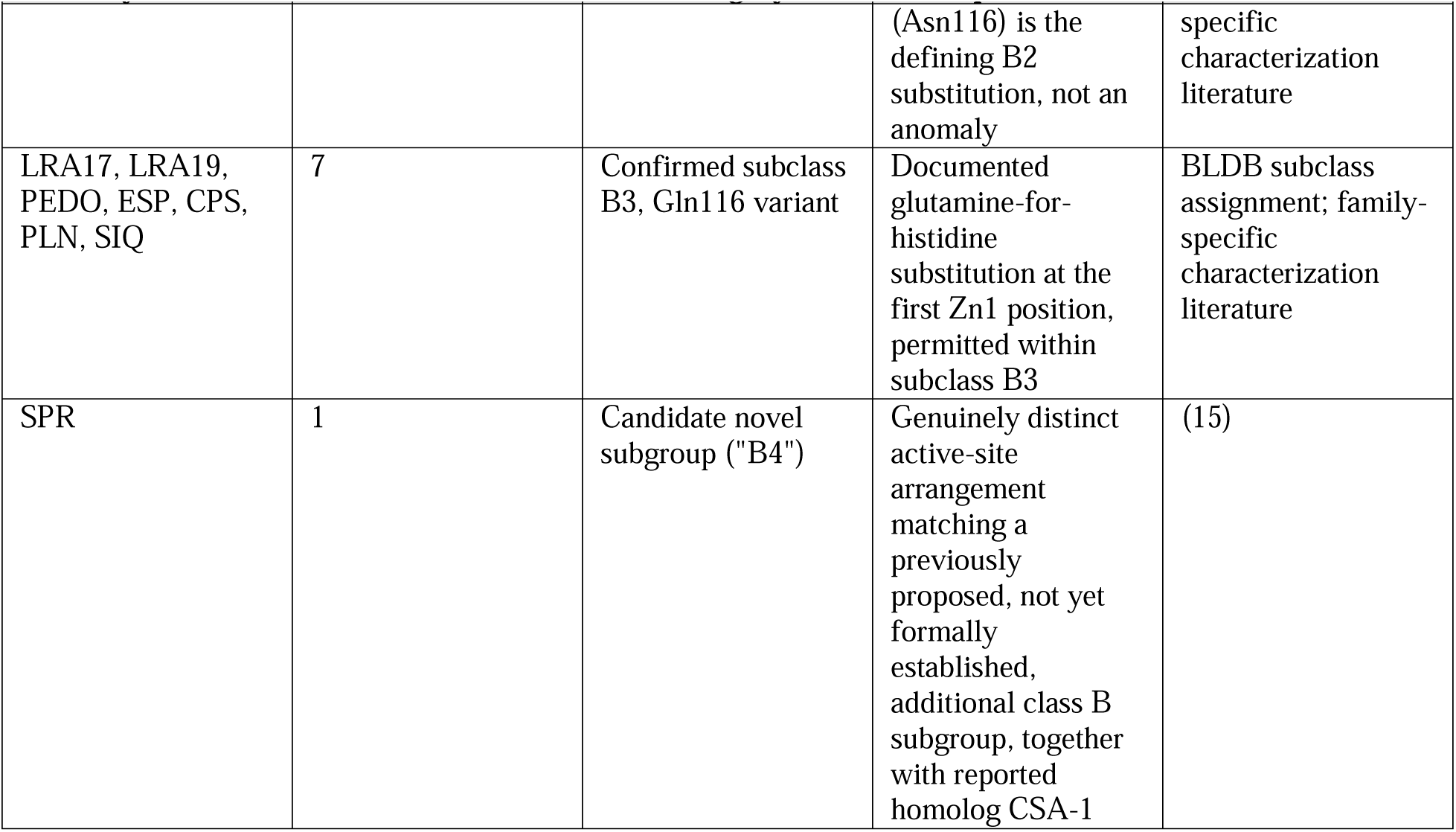
Resolution of 11 of the 16 target-set families lacking an exact canonical zinc-ligand motif match. SER and SSE (subclass B3 by BLDB and (20)) match neither HxHxD nor Q/N-x-H-x-D and are not explained by the Gln116 substitution; VMH, AHM and CrxA are treated in Table 5.

### Two independent numbering-transfer methods identify a small number of families with genuinely divergent active-site structure

For most of the 87 target families, BBL zinc-ligand numbering could be transferred with confidence directly from sequence alignment to the appropriate subclass reference enzyme (Needleman–Wunsch global alignment, BLOSUM62 substitution matrix, affine gap penalties) (21, 22), giving a clean, monotonic, fully mapped correspondence for the six canonical zinc-ligand positions. Five families, however, produced non-monotonic, gap-riddled or otherwise flagged-unreliable sequence alignments against every available reference enzyme of the expected subclass and could not be confidently anchored by sequence alone: BB13, VMH, BB32, AHM, and CrxA. For these five, we additionally performed structure-based residue transfer, superimposing each family’s predicted structure directly onto the most appropriate available historical-benchmark reference structure in PyMOL and reading the corresponding zinc-ligand positions from the resulting structural alignment (Table 5; Figure 4).

**Figure 4.**
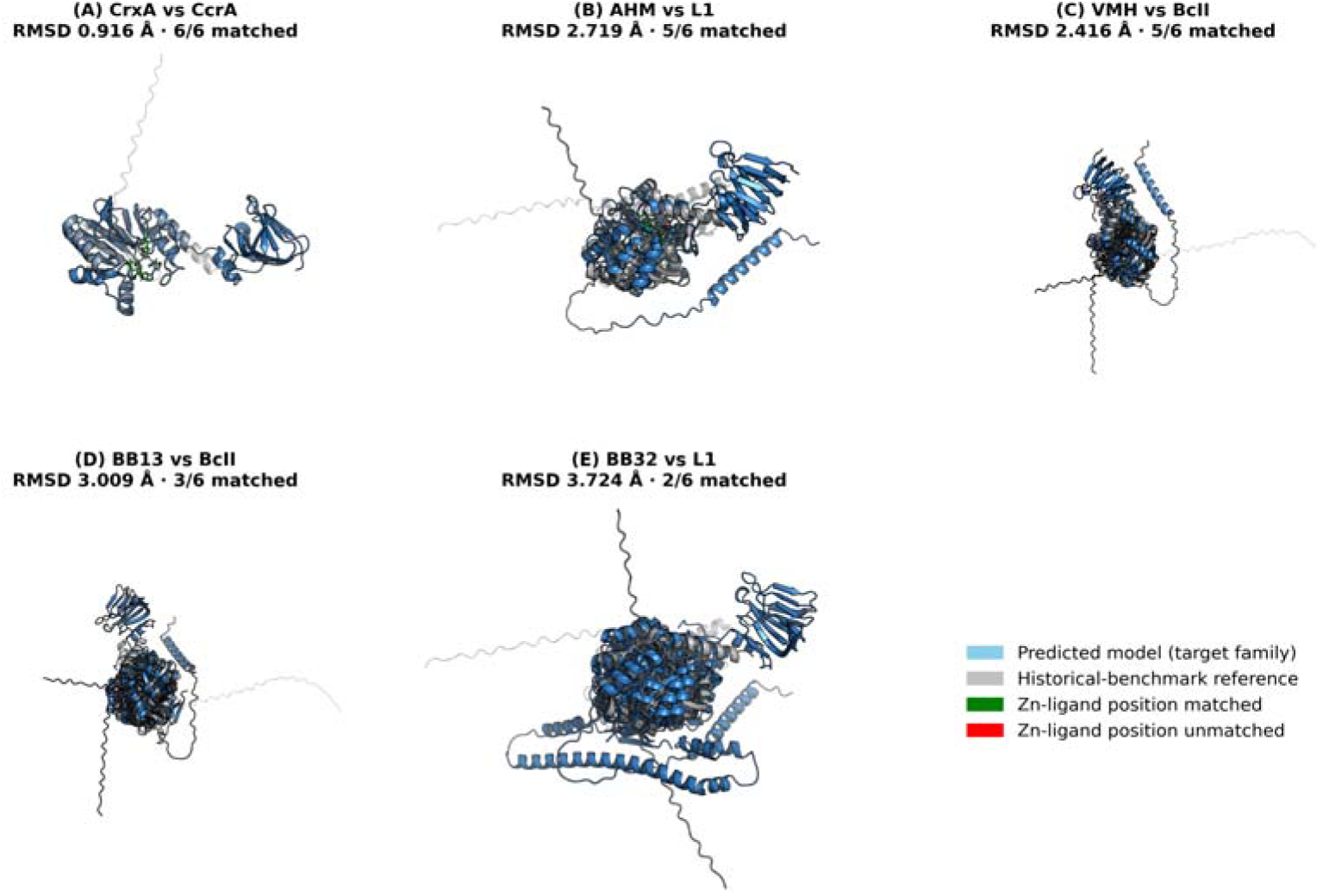
Structural superposition of the predicted models for CrxA, AHM, VMH, BB13, and BB32 (blue) onto their respective historical-benchmark reference structures (gray; CcrA, L1, BcII, BcII, and L1, respectively) in PyMOL. Panel labels correspond to Table 5: (A) CrxA vs. CcrA, RMSD 0.916 Å, 6/6 zinc-ligand positions matched; (B) AHM vs. L1, RMSD 2.719 Å, 5/6 positions matched; (C) VMH vs. BcII, RMSD 2.416 Å, 5/6 positions matched; (D) BB13 vs. BcII, RMSD 3.009 Å, 3/6 positions matched; (E) BB32 vs. L1, RMSD 3.724 Å, 2/6 positions matched. Green sticks indicate matched zinc-ligand positions; red sticks indicate positions that could not be matched by structural superposition (none for CrxA, one position each for AHM and VMH; three positions for BB13 and four for BB32; see Table 5 for the full per-position breakdown for all five families).

**Figure 5.**
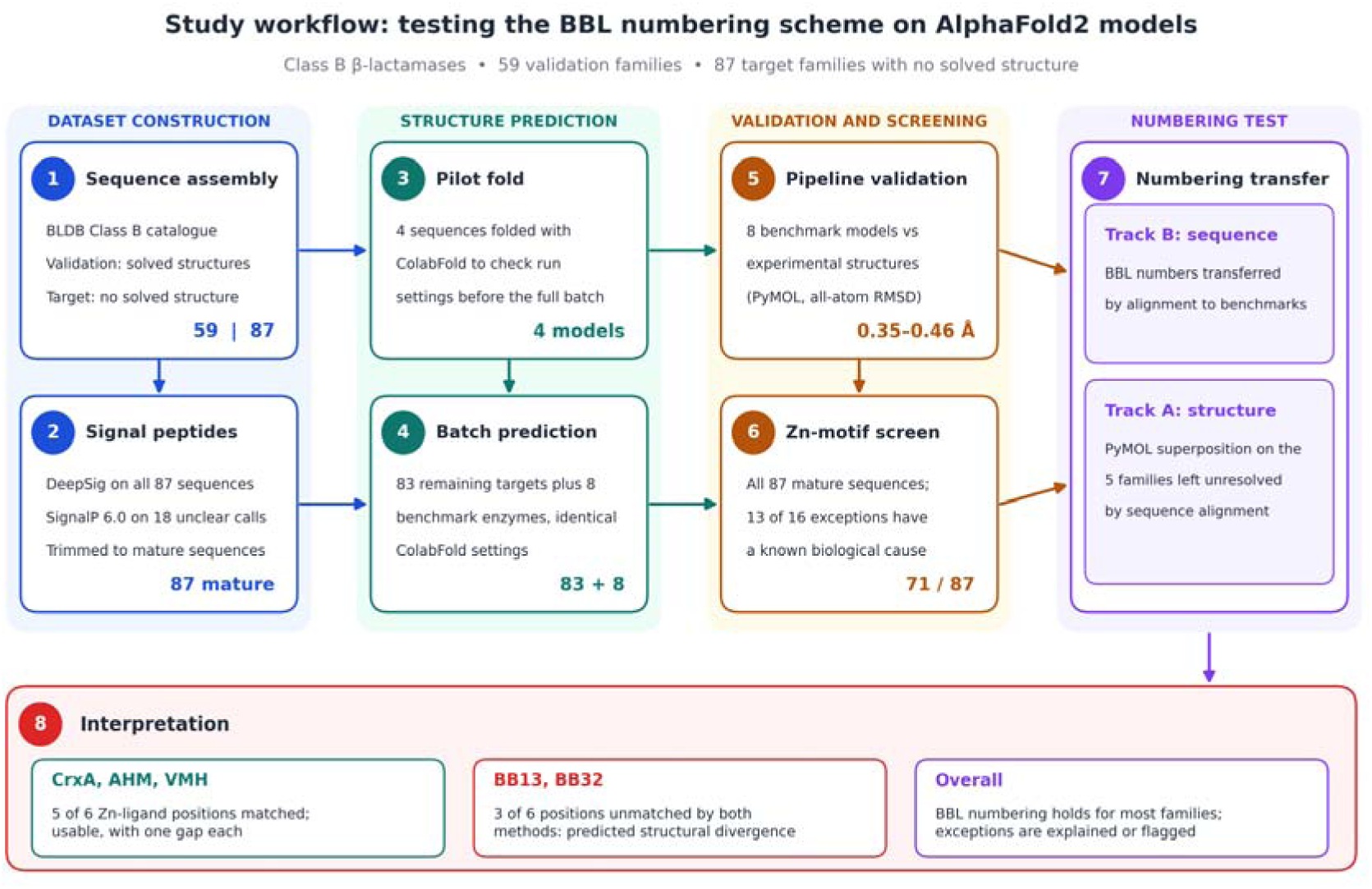
Schematic overview of the study workflow: BLDB-derived dataset construction, signal-peptide processing, ColabFold structure prediction, pipeline validation against the eight-enzyme historical benchmark, and paired sequence- and structure-based BBL numbering transfer to the 87-family target set.

**Table 5.** Structure-based zinc-ligand numbering transfer for the five families not resolved by sequence alignment alone.

| Family | Reference structure | RMSD (Å) | Zn1 result (3 positions) | Zn2 result (3 positions) | Verdict |
| --- | --- | --- | --- | --- | --- |
| CrxA | CcrA | 0.916 | Arg (subst.), His, His | Asp, Asn (subst.), His | Resolved; all six positions mapped; Arg substitution also found by sequence-based method |
| AHM | L1 | 2.719 | His, Asp (subst.), [gap] | Asp, His, His | Usable; one unmatched position |
| VMH | BcII | 2.416 | Gly (subst.), Asp (subst.), [gap] | Asp, Val (subst.), His | Mapped at 5/6 positions but three non-canonical residues; provisional |
| BB13 | BcII | 3.009 | His, His, [gap] | Asp, [gap], [gap] | Unresolved; 3/6 positions unmatched despite 81% chain coverage |
| BB32 | L1 | 3.724 | His, [gap], [gap] | [gap], Pro (subst.), [gap] | Unresolved; 4/6 positions unmatched despite 74% chain coverage |

One of the five (CrxA) was fully resolved by this structural approach: superposition onto CcrA (RMSD 0.916 Å) gave a clean, monotonic correspondence for all six zinc-ligand positions, with an arginine substituted for the expected first Zn1 histidine and an asparagine at the position of the Zn2 cysteine — the arginine substitution matched, independently, the result separately obtained from CrxA’s sequence-based alignment (the sequence alignment left the Zn2 middle position as a gap) once that alignment was re-anchored to CcrA rather than to the default subclass B1 reference (BcII), consistent with CrxA’s published closer phylogenetic relationship to the CcrA/CfiA lineage than to BcII (16). Two further families (AHM, a previously characterized Clostridioides difficile enzyme with a non-canonical active site (17), and VMH, a Vibrio vulnificus MBL-fold metallohydrolase originally reported to show no significant sequence identity to experimentally confirmed class B β-lactamases but independently classified as subclass B1 in BLDB (18)) were mapped at five of six positions each, at a moderately elevated RMSD to their respective reference structures (2.7 Å and 2.4 Å, respectively). AHM carries one Asp-for-His substitution at the second Zn1 position and was resolved as usable; VMH carries non-canonical residues at three of its five mapped positions (Gly and Asp at Zn1, Val at the Zn2 cysteine position) and is treated as provisional.

The remaining two families, BB13 and BB32, could not be confidently resolved for precise zinc-ligand numbering by either method. Structural superposition gave elevated RMSDs (3.0 Å and 3.7 Å, respectively) with three of six zinc-ligand positions unmatched for BB13 and four of six for BB32. To distinguish genuine structural divergence from a simple registration failure of the global superposition, we examined the fraction of each chain actually included in the structural alignment: 81% of BB13’s chain and 74% of BB32’s chain aligned to their reference structures, ruling out a small mis-anchored fragment (which would be expected to align only 30–50 residues) as the explanation. Both families retain an intact canonical zinc-ligand sequence motif (motif screen; zn_motif_prescreen.csv), indicating that the divergence is confined specifically to the three-dimensional arrangement of the active-site loop region rather than reflecting loss of the catalytic residues themselves. We interpret this as evidence of predicted structural divergence at the active site in these two families, which experimental structure determination would be needed to confirm — the kind of finding that a purely sequence-based renumbering approach, however well designed, cannot by construction flag, since it does not evaluate three-dimensional structure at all.

### The subclass-diagnostic active-site loop (BBL 61–65) is confirmed present in B1 and absent in B3

Galleni et al. describe a loop closing the active site of subclass B1 enzymes, spanning BBL residues 61–65, as present in subclass B1, absent in subclass B3, and probably absent in subclass B2 (6). Direct inspection of the primary reference alignment (6, 7) confirms that this loop corresponds to the region between conserved β-strands S2 and S3 (labeled L1 in the original figures), and that all five subclass B1 historical-benchmark structures carry genuine residues at this position while both subclass B3 historical-benchmark structures (L1, FEZ-1) show a contiguous alignment gap at the equivalent position — a direct, unambiguous confirmation of the published subclass distinction. A sequence-length proxy analysis attempting to test this feature computationally across all 87 target-set families (residue count from the sequence start to each family’s own Zn1 motif, as a stand-in for loop length) proved confounded by unrelated indels elsewhere in the same N-terminal region — precisely the kind of alignment ambiguity the BBL scheme exists to resolve — and produced no clean subclass separation (B1 mean 81.8 residues to the Zn1 motif versus B3 mean 88.5 residues (n = 35; B1, n = 44), the opposite direction from the length difference the loop’s presence/absence alone would predict). We therefore report the loop’s subclass distribution as confirmed at the level of the primary reference structures, as is standard practice when citing an established structural feature of a reference scheme, rather than claiming an independent per-family re-derivation for the target set; this is stated explicitly as a scope limitation in the Discussion.

## DISCUSSION

More than twenty years after the last update to the standard class B β-lactamase numbering scheme, and using a structure-prediction technology unavailable at the time either version was proposed, we find that the BBL scheme’s core zinc-ligand correspondences generalize well to the much larger and more diverse body of class B sequence now known. Of the 87 never-crystallized families examined here, the canonical zinc-ligand motif transferred exactly in 82%, and 11 of the 16 remaining exceptions had a specific, literature-supported biological explanation rather than representing a failure of the numbering scheme or the prediction pipeline (the other five, SER, SSE, VMH, AHM, and CrxA, are addressed below; SER and SSE are B3 enzymes by BLDB with genuine deviations from the motif (Glu at the first Zn1 position in SSE; Arg at the second Zn1 position in SER)): two categories of exception (three confirmed subclass B2 members and seven confirmed subclass B3 members) reflect substitutions the scheme’s own subclass-specific consensus already anticipates, and a further exception (SPR) corresponds to a previously proposed, though still only tentatively established, additional class B subgroup (15). Independent structural superposition on the small number of families whose sequence-level anchoring was most ambiguous reinforced rather than contradicted this picture: two of five (AHM and CrxA) were resolved as usable, CrxA by both sequence and structure methods, which agree on the Zn1 arginine substitution once anchored to its correct phylogenetic reference; VMH was mapped but remains provisional, with non-canonical residues at three positions; and only two families (BB13 and BB32) showed a signature of predicted active-site structural divergence, being flagged as unreliable by the sequence method and only partly mapped by structural superposition.

These two families are, in our view, the most interesting positive finding of this study rather than a limitation of it. Both retain the canonical zinc-ligand sequence motif, ruling out simple loss of the catalytic machinery, yet both show substantially elevated structural RMSD to every available subclass reference and half or more of their six mapped zinc-ligand positions unresolved (three of six for BB13 and four of six for BB32) by direct structural superposition, despite roughly three-quarters or more of each chain aligning well overall. This is precisely the class of finding that a sequence-only renumbering approach — however well constructed — cannot surface, because it requires evaluating the three-dimensional consequences of sequence divergence rather than the sequence divergence itself. Oelschlaeger et al. have separately identified the broader class B numbering problem this study addresses and describe an in-development tool for sequence-based renumbering via profile hidden Markov models (10); that approach and the present one are complementary rather than competing, since sequence-based renumbering can scale to far more sequences at far lower computational cost, while the structure-prediction approach used here is needed specifically to detect the kind of local active-site divergence identified in BB13 and BB32. We suggest that BB13 and BB32 (and, more provisionally, the candidate B4-type family SPR) are strong candidates for direct experimental structural characterization to confirm and further resolve the nature of their active-site divergence.

This study has several limitations that should inform how its conclusions are used. First, subclass B2 representation in the target set is limited to three families (PFM, CVI, YEM), reflecting the genuine rarity of subclass B2 among named class B enzymes generally (only five families are recorded in BLDB for this subclass in total) rather than an artifact of our filtering; conclusions about subclass B2 specifically should be treated as more provisional than those for B1 and B3 pending characterization of additional B2 representatives as they are described. Second, five target-set sequences (PAN, SFB, SLB, BB32, and AHM) have no confident signal-peptide call and were retained at full length rather than trimmed on a low-confidence prediction (three of them, PAN, SFB, and SLB, were genuinely ambiguous under SignalP); because their catalytic-domain confidence and zinc-ligand numbering were unaffected, this does not compromise the numbering results reported here, but it is flagged for resolution should these families’ mature boundaries become independently known. Third, we did not attempt to extend the present analysis from the six canonical zinc-ligand positions to the full set of fourteen structurally conserved sequence fragments identified by Garau et al. (7), since extracting precise residue boundaries for all fourteen fragments from the original alignment figures proved too error-prone to justify given that (i) the active-site positions most relevant to catalysis and substrate specificity are already covered by the present analysis, and (ii) a preliminary sequence-based attempt at this extension was confounded by the same unrelated indel variation described above for the active-site loop check. We regard this as a deliberate scope decision rather than an unexamined gap, and note it explicitly so that future work extending the present dataset to the full fragment set can build directly on the sequence and structure resources described here.

The JOHN family sequence used here matches the original characterization of the Flavobacterium johnsoniae carbapenemase (19) exactly at the residue level, providing an independent, literature-anchored check on this portion of the sequence-preparation pipeline.

Taken together, these results support extending the validity of the BBL numbering scheme into structural space that did not exist, computationally, at the time the scheme was proposed, and they illustrate a general template — pipeline validation against a small experimental benchmark, followed by paired sequence- and structure-based residue-transfer analysis of a much larger predicted-structure set, with explicit resolution or transparent reporting of every resulting exception — for extending structure-dependent numbering and classification schemes across any protein family for which structural characterization has lagged behind sequence discovery.

## MATERIALS AND METHODS

### Database sources and dataset construction

Class B β-lactamase sequences, subclass assignments, and structural-availability annotations were drawn from the Beta-Lactamase DataBase (BLDB) (8). The full class B catalogue (1,021 named variants across 129 family–subclass entries) was filtered to families for which no member variant had a recorded solved structure, yielding 96 initial candidate family–subclass entries (95 unique families). This candidate list was cross-checked against the 37 protein families of the validation set of confirmed solved-structure enzymes to identify BLDB annotation errors, spot-checked against RCSB PDB and the primary literature for a representative subsample, and independently re-verified in a further literature pass; corrections at each stage (detailed in Results) yielded a final target set of 87 families with no solved structure. A parallel validation set of solved-structure enzymes (59 unique sequences after removal of one duplicate fragment and addition of VIM-1, independently representing 58 of 60 unique enzymes drawn from 37 BLDB protein families and 590 individual PDB entries) was assembled from the same catalogue by UniProt ID mapping (54 of the 59 enzymes with an identifiable accession) supplemented by NCBI Batch Entrez retrieval using BLDB-listed GenPept identifiers for the remaining five (CHI-1, CSR-1, RM3, MYO-1, PNGM-1). An eight-enzyme historical benchmark subset (BcII, CcrA, IMP-1, VIM-2, BlaB, CphA, L1, FEZ-1), corresponding to the structures used in the original 2004 BBL update (7), was selected from the validation set for pipeline validation and structural anchoring.

### Signal peptide identification and removal

Target-set sequences were screened for N-terminal signal peptides using DeepSig (11). Sequences for which DeepSig did not call a signal peptide were additionally scanned for N-terminal hydrophobicity using the Kyte–Doolittle scale and, where a hydrophobic-core signature was present despite the negative DeepSig call, submitted to SignalP 6.0 (organism setting "Other," slow/sensitive mode) as an independent second prediction (12). A signal peptide was trimmed only where a specific cleavage site was predicted with posterior probability ≥ 0.70; sequences with no confident call under either method were retained at full length and flagged as a stated limitation. Validation-set and historical-benchmark sequences were not subjected to signal-peptide trimming, since these enzymes’ mature-sequence boundaries and experimental structures are independently known or, for the historical benchmark, were separately confirmed by SignalP 6.0 as described in Results.

### Structure prediction

All structure predictions were generated using ColabFold v1.6.3, an implementation of AlphaFold2 using MMseqs2 for multiple-sequence-alignment generation (13, 14). Settings were held identical across every prediction job: no structural templates (template_mode = none), automatic model-type selection, three recycles, paired/unpaired MMseqs2 UniRef and environmental-database MSA generation, and Amber relaxation of the top-ranked output model (num_relax = 5). Each sequence was submitted as a single, non-complexed chain. Per-model quality was assessed from the ColabFold-reported per-residue predicted local distance difference test (pLDDT) score and predicted TM-score (pTM); a first-30-residue mean pLDDT was additionally computed for each model as an indicator of residual N-terminal disorder consistent with an unresolved, uncleaved signal peptide.

### Validation against experimental structures

Predicted structures for the eight-enzyme historical benchmark set were superimposed on their corresponding experimental structures using the PyMOL align command, which performs an initial sequence-based alignment followed by iterative structural refinement with outlier rejection. Experimental reference structures were PDB entries 1BC2 (BcII), 1ZNB (CcrA), 1DDK (IMP-1, apo form), 1KO3 (VIM-2), 1M2X (BlaB), 1X8G (CphA, wild-type native form), 1SML chain A (L1), and 5WCK (FEZ-1, the current RCSB-designated replacement for the original 1K07 deposition). Non-protein heteroatoms other than the catalytic zinc ions were removed from experimental structures prior to alignment. Reported RMSD values were calculated over all atoms (not Cα-only), using PyMOL’s default five cycles of iterative outlier rejection. Because rejection cycles progressively exclude poorly superposing atoms—typically at flexible termini or loop regions—to isolate a well-conserved structural core, the number of atoms retained in the final alignment is substantially smaller than each model’s total atom count (e.g., BcII: 1395 of 4025 total atoms; Table 2); this is the intended behavior of the algorithm rather than an artifact. As a robustness check, we additionally computed Cα-only RMSD without outlier rejection (0 cycles) for all eight enzymes; this produced markedly unstable values for several (e.g., BcII 14.9 Å, L1 4.7 Å), consistent with a small number of poorly conserved residues—most likely within the flexible, unresolved signal-peptide region discussed above— dominating a naive superposition, and confirming that outlier rejection is necessary rather than a means of inflating apparent accuracy.

### Sequence-based residue-numbering transfer

BBL zinc-ligand numbering was transferred from the historical benchmark reference enzymes to each of the 87 target-set sequences by global pairwise alignment (Needleman–Wunsch algorithm, BLOSUM62 substitution matrix, affine gap penalties: gap opening −11, gap extension −1) (21, 22), implemented directly rather than via a third-party alignment package. For each reference enzyme, the offset between its precursor (signal-peptide-containing) sequence numbering, as deposited, and its published BBL mature-sequence numbering (6, 7) was determined empirically by identifying the single offset value that correctly reproduced the expected amino acid identity at all six canonical zinc-ligand positions simultaneously; this offset was then used to translate alignment-derived positions into BBL-consistent mature-sequence numbering. Each target family was aligned against the reference enzyme(s) of its assigned subclass (BcII or CcrA for subclass B1; CphA for subclass B2; L1 or FEZ-1 for subclass B3); where an initial alignment against the default subclass reference produced a non-monotonic or heavily gap-riddled correspondence at the zinc-ligand positions, an alternative, phylogenetically more appropriate reference enzyme from the same subclass was tried before a family was classified as unresolved by sequence alignment alone.

### Structure-based residue-numbering transfer

For target-set families whose zinc-ligand numbering could not be confidently resolved by sequence alignment (defined as producing a non-monotonic or gap-riddled correspondence at the zinc-ligand positions against every available subclass reference), structure-based numbering transfer was performed by superimposing the family’s predicted structure directly onto the most phylogenetically appropriate available historical-benchmark reference structure in PyMOL (cmd.super), extracting the resulting residue-level structural alignment (cmd.get_raw_alignment), and mapping each reference enzyme’s known BBL zinc-ligand position onto the corresponding target-structure residue. Alignment coverage was additionally assessed as the fraction of each target chain’s Cα atoms included in the structural alignment, to distinguish genuine local structural divergence at the active site (high overall coverage, localized non-correspondence at specific ligand positions) from a global registration failure (low overall coverage).

### Sequence-motif screening for zinc-ligand conservation

All 87 mature target-set sequences were scanned computationally for the canonical zinc-ligand motif spanning the Zn1 histidine pair (or subclass-appropriate substitute) and the Zn2 aspartate, using the regular-expression patterns HxHxD (subclass B1 and some B3 members), Q/N-x-H-x-D (subclass B2 and Gln116-variant B3 members (SER and SSE lack this pattern)), and H-x-D-x-D-H (observed in specific non-canonical B3-like enzymes), calibrated against the known mature-numbering positions of the historical benchmark enzymes (Table 1). Sixteen families lacked an exact HxHxD match: ten of these matched Q/N-x-H-x-D, one (AHM) matched H-x-D-x-D-H, and five (SPR, SER, SSE, VMH, CrxA) matched none of the three patterns. These families were individually resolved by cross-reference to BLDB subclass assignment and the primary characterization literature for that family. The sequence-alignment method additionally flagged BB11, which has an exact HxHxD match, as carrying Asp rather than His at the BBL 196-equivalent Zn1 position (Asp153 in the mature sequence).

### Active-site loop verification

Presence or absence of the subclass-diagnostic active-site-closing loop (BBL residues 61–65) was assessed by direct inspection of the published reference alignment (6, 7) for the historical benchmark structures, and, as a secondary computational check, by comparing the residue count from each target-set sequence’s start to its own Zn1 motif position as a proxy for loop length, stratified by BLDB subclass assignment.

### Statistical and computational environment

Sequence parsing, motif scanning, and summary statistics were implemented in Python. Multiple-sequence and pairwise alignment (excluding the from-scratch Needleman–Wunsch implementation described above) and structural manipulation were performed using PyMOL (v3.2.0a for local structural work; Schrödinger, LLC). No commercial or proprietary sequence databases were used at any stage.

## DATA AVAILABILITY

The 87-family target-set mature-sequence FASTA file, the 59-sequence validation-set FASTA file, the eight-enzyme historical-benchmark reference FASTA file, the BLDB-derived validation-set and subclass-assignment spreadsheets, the signal-peptide screening audit table, per-family ColabFold confidence scores (pLDDT, PAE, pTM) for all 87 target families, the zinc-motif screen, the sequence-alignment (Track B) and structure-based (Track A) zinc-ligand mapping outputs, and the benchmark RMSD and score tables have been deposited in a public repository. Zenodo, https://doi.org/10.5281/zenodo.23087005.

## ACKNOWLEDGMENTS

This research received no specific grant from any funding agency in the public, commercial, or not-for-profit sectors. All structure predictions were run in Google Colaboratory using ColabFold v1.6.3; downstream analysis was performed locally. No competing interests are declared. Author contributions (CRediT): M.M.K.N.: Conceptualization, Methodology, Software, Formal analysis, Investigation, Data curation, Writing – original draft, Writing – review & editing, Visualization. H.A.K.: Supervision, Validation, Writing – review & editing.

## Notes

https://doi.org/10.5281/zenodo.23087005

### Competing Interest Statement

The authors have declared no competing interest.

